# Bacterial communities of herbivores and pollinators that have co-evolved *Cucurbita* spp

**DOI:** 10.1101/691378

**Authors:** Lori R. Shapiro, Madison Youngblom, Erin D. Scully, Jorge Rocha, Joseph Nathaniel Paulson, Vanja Klepac-Ceraj, Angélica Cibrián-Jaramillo, Margarita M. López-Uribe

## Abstract

Insects, like all animals, are exposed to diverse environmental microbes throughout their life cycle. Yet, we know little about variation in the microbial communities associated with the majority of wild, unmanaged insect species. Here, we use a 16S rRNA gene metabarcoding approach to characterize temporal and geographic variation in the gut bacterial communities of herbivores (*Acalymma vittatum* and *A. trivittatum*) and pollinators (*Eucera (Peponapis) pruinosa*) that have co-evolved with the plant genus *Cucurbita* (pumpkin, squash, zucchini and gourds). Overall, we find high variability in the composition of bacterial communities in squash bees and beetles collected from different geographic locations and different time points throughout a growing season. Still, many of the most common OTUs are shared in *E. (P*.*) pruinosa, A. vittatum* and *A. trivittatum*. This suggests these insects may be exposed to similar environmental microbial sources while foraging on the same genus of host plants, and that similar microbial taxa may aid in digestion of *Cucurbita* plant material. The striped cucumber beetle *A. vittatum* can also transmit *Erwinia tracheiphila*, the causal agent of bacterial wilt of cucurbits. We find that few field-collected *A. vittatum* individuals have detectable *E. tracheiphila*, and when this plant pathogen is detected, it comprises less than 1% of the gut bacterial community. Together, these results are consistent with previous studies showing that plant feeding insects have highly variable gut bacterial communities, and provides a first step towards understanding the spatiotemporal variation in the microbial communities associated with herbivores and pollinators that depend on *Cucurbita* host plants.

## Introduction

Herbivorous insects that consume leaf, floral, stem or root tissues are exposed to diverse communities of microbes associated with plants and/or soil. Many microbes ingested by insects likely pass transiently through the digestive tract and may not confer any impact on the host insect (Oliver et al. 2008, Hammer et al. 2017, Hammer et al. 2019). However, some microbes are able to colonize insects as hosts, and can have diverse and consequential – yet often cryptic – roles in mediating ecological interactions between plants, other insects and other microbes (Shapiro et al. 2012, Chung et al. 2013, Shapiro et al. 2013, Wang et al. 2016, Mason et al.2018). For example, many pollinators likely transfer microbes between flowers during normal foraging that may directly affect other visiting insects, or indirectly alter traits determining floral attractiveness (Yang et al., McArt et al. 2014, Ravoet et al. 2014, Vannette and Fukami 2016, Rering et al. 2018, Figueroa et al. 2019). Some insect pollinators and herbivores encounter plant pathogens while feeding, and can then transmit these pathogens from infected to healthy plants (Raguso and Roy 1998, Alexandrova et al. 2002, López-Villavicencio et al. 2007, Degnan et al. 2009, Shapiro et al. 2012, Shapiro et al. 2014, Mauck et al. 2018, Shapiro and Mauck 2018, Cellini et al. 2019).Yet, for almost all insect taxa, we still lack an understanding of the spatial and temporal diversity of microbes that insects are naturally exposed to, which microbes are able to persistently colonize insects as hosts vs. transiently pass through the digestive tract, and the impacts of different microbes on ecological interactions.

The pollinators and herbivores that have co-evolved with the plant genus *Cucurbita* (Cucurbitaceae) are an ideal system to investigate how a shared host plant affects microbiome composition for insect species that have different life histories and ecological functions. *Cucurbita* (zucchini, squash, pumpkin, and some yellow-flowered gourds) is comprised of 14 closely related species that are native to the New World tropics and subtropics (Whitaker and Bemis 1975, Metcalf and Lampman 1989, Kates et al. 2017, Castellanos-Morales et al. 2018). All bee species in the genus *Eucera* and sub-genera *Peponapis* and *Xenoglossa* (Hymenoptera: Anthophila: Apidae: Eucerini: *Eucera*) – commonly known as squash bees – are obligate specialists of *Cucurbita* pollen (Baker and Hurd Jr 1968, Hurd et al. 1971, Hurd et al. 1974). The emergence of squash bees in the summer coincides with the *Cucurbita* bloom period, and these ground-nesting, solitary bees exclusively use pollen from *Cucurbita* flowers to feed their brood (Hurd and Linsley 1964, Hurd and Linsley 1967, Hurd et al. 1971, Hurd et al. 1974). *Acalymma* (Barber) leaf beetle herbivores (Coleoptera: Chrysomelidae: Luperini: Diabriticina) have also co-evolved with *Cucurbita*, and all ∼70 species of *Acalymma* are obligately dependent on *Cucurbita* in all life stages. *Acalymma* larvae are rootworms, and adults feed on above-ground leaves, flowers and fruits (Barber 1946, Metcalf et al. 1980, Munroe and Smith 1980, Ferguson and Metcalf 1985, Ferguson et al. 1985, Tallamy and Krischik 1989, Tallamy and Gorski 1997, Tallamy et al. 1997, Kistler et al. 2015).

The squash bee (*Eucera (P*.*) pruinosa*) and the striped cucumber beetle (*Acalymma vittatum*) are the two most common *Cucurbita-*associated insects in temperate Eastern North America. *E. pruinosa* has recently expanded its geographic range into this region by following the human-mediated dispersal of domesticated *Cucurbita pepo* from the subtropical Eastern United States (Petersen and Sidell 1996, Monaghan et al. 2006, Smith 2006, López-Uribe et al. 2016), and it is likely that *A. vittatum* has a similar demographic history. In temperate Eastern North America, both *E. pruinosa* and *A. vittatum* exclusively rely on domesticated plants grown in human managed agro-ecosystems as a food resource. Temperate Eastern North America is also the only region worldwide where *A. vittatum* transmits *Erwinia tracheiphila* Smith (Enterobacteriaceae), the causal agent of bacterial wilt of cucurbits, which causes millions of dollars annually in direct yield losses and indirect preventative measures (Shapiro et al. 2014, Shapiro et al. 2016, Shapiro et al. 2018b). *E. tracheiphila* can persistently colonize the digestive tract of beetle vectors after beetles feed on the foliage of wilting, symptomatic plants (Rand and Enlows 1916, Rand and Cash 1920, Rand and Enlows 1920, Smith 1920, Shapiro et al. 2012, Shapiro et al. 2014). However, dynamics of vector colonization by *E. tracheiphila* in the field, and potential interactions with other microbes in the beetle’s digestive tract remain almost completely unknown (Fleischer et al. 1999, Shapiro et al. 2014).

Here, we use 16S rRNA gene barcode sequencing to describe variation in the bacterial communities of the squash bee *Eucera (P*.*) pruinosa*, and the striped cucumber beetles *Acalymma vittatum* and *A. trivittatum*. Insects were field-collected from their native range in the United States Southwest and in Central Mexico, and from their introduced range in temperate North America. We find high overall variation in bacterial community composition within and between beetles and bees that specialize on *Cucurbita*, but substantial overlap in the most common bacterial OTUs. We also find that few *Acalymma* beetles are colonized by detectable *Erwinia tracheiphila*, and that OTUs that are the closest match to *E. tracheiphila* are a minor component of the beetle vector’s gut bacterial community. Together, these results suggest the shared *Cucurbita* host plants are likely a strong factor influencing microbial community composition of bees and beetles that have co-evolved with these hosts.

## Methods

### The study system

Current taxonomic assignments recognize 14 closely related *Cucurbita* species (Kates et al. 2017). All are fast-growing vines, and many wild species are common throughout the New World tropics and subtropics (Kates et al. 2017, Castellanos-Morales et al. 2018). All *Cucurbita* produce large, yellow, sweetly scented flowers that only open a single morning from sunrise until midday, and then dehisce. Flowers release a blend of volatile organic compounds that act as long distance host location cues for co-evolved pollinators and herbivores. *Cucurbita* also produce a group of oxygenated tetracyclic tri-terpenes called ‘cucurbitacins’ that are among the most bitter compounds known, with broad-spectrum toxicity against almost all insect and mammalian herbivores (David and Vallance 1955, Fraenkel 1959, Whitaker and Bemis 1964, Chambliss and Jones 1966, Shang et al. 2014, Zhou et al. 2016). For Diabroticite leaf beetle herbivores (Coleoptera: Chrysomelidae: Luperini: Diabroticina), cucurbitacins are arrestants and feeding stimulants. All *Acalymma* are specialists on *Cucurbita* in all life stages, while *Diabrotica* and *Ceratoma* spp. only visit *Cucurbita* pharmacophagically as adults to sequester cucurbitacins (Metcalf et al. 1980, Tallamy and Krischik 1989, Dinan et al. 1997, Tallamy et al. 1997, Tallamy 1998). *Acalymma* larvae feed underground on *Cucurbita* roots, and adults feed on leaves, flowers and fruits (Barber 1946, Munroe and Smith 1980, Tallamy 1999, Eben and Espinosa 2013, Shapiro and Mauck 2018).

While insect herbivory has driven the evolution of *Cucurbita* functional traits over their millions of years of co-evolutionary history, humans have shaped the more recent evolution of many *Cucurbita* species. *Cucurbita pepo* was the first plant domesticated for agriculture in the Americas, and *Cucurbita moschata* was the second (Smith 1993, Smith 1997, Piperno and Stothert 2003, Smith 2006). As the glaciers retreated during the Holocene around 5,000 years ago, cultivated *Cucurbita* were the first crop plant introduced into temperate Eastern North America by humans for agriculture from a second independent domestication of *Cucurbita pepo* in the Mississippi River Valley (Petersen and Sidell 1996, Monaghan et al. 2006, Smith 2006). It is likely that movement of domesticated *Cucurbita pepo* varieties into temperate Eastern North America had profound – but still largely unknown – impacts on the biotic community that had co-evolved with *Cucurbita* for millions of years. *Eucera (P*.*) pruinosa* (López-Uribe et al. 2016) was the only *Cucurbita* specialist pollinator species, and *Acalymma vittatum* the only obligate herbivore species that were able to colonize the new temperate geographic areas where humans introduced *Cucurbita* cultivars (Shapiro et al. 2018b). In the temperate Eastern United States, *Acalymma vittatum* can transmit the virulent bacterial pathogen, *Erwinia tracheiphila*. This pathogen does not occur anywhere else in the world outside of this limited geographic area, and inferences from *E. tracheiphila* genomics studies suggest it was human-mediated changes to cucurbit agro-ecosystems that recently drove its emergence in this region (Shapiro et al. 2015, Shapiro et al. 2016, Andrade-Domínguez et al. 2018, Shapiro et al. 2018a, Shapiro et al. 2018b).

### Insect collection

Insects were field collected from flowers of healthy (*ie*, not infected with *Erwinia tracheiphila*) wild and cultivated *Cucurbita* spp. host plants, and all insects per location came from the same field or wild plant population (Table 1). Insects were killed in 95% ethanol and stored at −80C. Beetles were identified to species by LRS, and bees were identified to species by MMLU.

**Table 1:** Collection location, date, and host plant for all insects sequenced in this study

| Insect Species | Collection Location | Host Plant | Number of Insects Sequenced | Collection Date | Latitude | Longitude |
| --- | --- | --- | --- | --- | --- | --- |
| *Acalymma vittatum* | Christian Herter Community Garden, Watertown, Massachusetts | Cultivars of *Cucurbita* spp. in a diverse community garden | 6 | June 7, 2016 | 42.366338 | -71.134525 |
| *Acalymma vittatum* | Christian Herter Community Garden, Watertown, Massachusetts | Cultivars of *Cucurbita* spp. in a diverse community garden | 7 | July 12, 2016 | 42.366338 | -71.134525 |
| *Acalymma vittatum* | Christian Herter Community Garden, Watertown, Massachusetts | Cultivars of *Cucurbita* spp. in a diverse community garden | 6 | August 14, 2016 | 42.366338 | -71.134525 |
| *Acalymma vittatum* | University of Vermont Horticultural Farm, Burlington, Vermont | *Cucurbita pepo* (Zucchini) | 6 | August 8, 2015 | 44.430052 | -73.204011 |
| *Acalymma vittatum* | Iowa State University Horticultural Research Station, Ames Iowa | Cultivated *Cucurbita* spp. and *Cucumis* spp. cultivars | 6 | August 18, 2016 | 42.107812 | -93.589034 |
| *Acalymma vittatum* | Gulf Coast Research and Extension Center, Mobile, Alabama | Cultivars of *Cucurbita* spp. | 6 | June 3, 2015 | 30.547286 | -87.88255 |
| *Acalymma vittatum* | University of Kentucky Horticultural Research Farm, Lexington, Kentucky | Acorn squash plot (*Cucurbita pepo*) | 6 | August 9, 2016 | 37.976154 | -84.533393 |
| *Acalymma trivittatum* | New Mexico | Wild roadside *Cucurbita foetidissima* | 2 | July 2, 2015 | 32.5225 | -108.730833 |
| *Acalymma trivittatum* | University of Arizona Pima Outreach Center, Tucson, Arizona | Cultivars of *Cucurbita* spp. and *Cucumis sativus* in demonstration garden beds | 6 | July 7, 2015 | 32.282077 | -110.94240 |
| *Acalymma trivittatum* | Quertaro, Mexico | Wild *Cucurbita foetidissima* on roadside | 8 | July 28, 2016 | 20.6889 | -100.326 |
| *Peponapis pruinosa* | California | Flowers of cultivated *Cucurbita* | 7 | August 2, 2010 | 38.7678 | -122.0379 |
| *Peponapis pruinosa* | Pennsylvania | Flowers of cultivated *Cucurbita* | 4 | July 29, 2011 | 39.9461 | -77.3018 |

### DNA extraction and sequencing

To degrade surface DNA, ethanol-preserved insects were immersed in 10% CoveragePlus for 30sec and then rinsed in molecular grade water for 10s. After surface DNA degradation, the whole insect was pre-ground with a bleach treated plastic pestle, and homogenized with a bead beater at the highest settings for 1 min. DNA was extracted from ground whole insects using Zymo Microbiomics (Tustin, CA) extraction kit following manufacturer’s instructions. The extracted DNA in each sample was quantified with Qubit Broad Spectrum Kit (Thermofisher), and the concentration in all samples was standardized to 10 ng/µl to be used as PCR template. Each DNA sample was amplified in triplicate using the bacterial 16S primers F515 (5’-GTGCCAGCMGCCGCGGTAA-3’) and R806 (5’-GGACTACHVGGGTWTCTAAT-3’), specific for the V4 region of 16S rRNA. In addition to the region specific for 16S rRNA hybridization, these oligonucleotides contain adapter, primer pad and linker sequences for Illumina (Supplemental Table 5). One unique reverse primer was used for each sample, which also contained a distinctive 12 nucleotide GoLay barcode (Supplemental Table 5) (Caporaso et al. 2011). Each 25 µl PCR reaction contained 0.25 μl of Q5 High Fidelity polymerase (New England Biolabs), 0.5 μl of 10 mM dNTPs, 500 nM of each primer and 5 μl of 5X Q5 reaction buffer. The PCR cycle program was: denaturation at 98°C for 30 s, then 35 cycles of 98°C for 10 s, 60°C for 30 s, and 72°C for 20 s; finally, 72 □ C for 2 min. The PCR products were checked via agarose gel electrophoresis for the correct size PCR product as a visual quality control. The three triplicate PCR products per sample were then pooled, gel purified using a Gel Purification Kit (Zymo Research, Tustin CA), and quantified using Qubit Broad Range kit (Thermofisher). Equal quantities of the pooled triplicates from each sample were then combined. This final pool containing all libraries was prepared for Illumina MiSeq sequencing by diluting to 4 nM and then denatured by mixing 1:1 with 0.2 N NaOH, for a final concentration of 2 nM of DNA and 0.1 N NaOH. The pooled libraries were then mixed with 1 volume of 2 nM denatured PhiX Sequencing Control (Illumina). Three primers were used for sequencing: read 1 of 251 cycles (5’-TATGGTAATTGTGTGCCAGCMGCCGCGGTAA-3’), read 2 for 251 cycles (5’-AGTCAGTCAGCCGGACTACHVGGGTWTCTAAT-3’) and index (5’-ATTAGAWACCCBDGTAGTCCGGCTGACTGACT-3’). The MiSeq kit V2 for 500 cycles (Illumina) was used for sequencing, following the manufacturer’s instructions.

### Data pre-processing and statistical analyses

All parameters for all pre-processing steps in Qiime and all statistical analyses in R are available in the qiime.txt file at https://github.com/lshapiro31/cucurbit.insect.microbiome. Qiime 1.8 was used for intial data processing, to demultiplex raw sequencing data and join paired reads (Caporaso et al. 2010). Chimeric sequences were identified *de novo* and removed using usearch 6.1 (Edgar 2010b). PyNast (Caporaso et al. 2009) was used to align the representative 16S sequences, and Usearch (Edgar 2010b) was used to cluster unique *de novo* OTUs at 97% similarity (Edgar 2010a). Taxonomy was assigned with Uclust (Edgar 2010b) using the greengenes database as the reference (DeSantis et al. 2006, McDonald et al. 2012). The nucleotide sequences of the most prevalent and abundant OTUs were used as a query in the BLASTN (Altschul et al. 1990) web interface to assess whether the taxonomy assignments from greengenes is accurate. When BLASTN provided a more accurate taxonomic assignment, that assignment was added in a separate column to the OTUs in the taxonomy files (Supplemental Tables 1, 2, 3, 4).

All statistical analyses were carried out in the R statistical computing environment (Team 2015) following the general pipelines established in phyloseq (McMurdie and Holmes 2013). OTUs that had less than five total reads were removed. One *Acalymma* sample (#SampleID 153; Supplemental Table 4) had only 2 total reads and was excluded from all analyses due to low sequencing coverage. OTUs that were assigned as chloroplast or mitochondria were filtered out of the OTU table, and *Wolbachia* OTUs were filtered to a new biom table to be analyzed separately. Phyloseq was used to rarefy the number of all reads per sample to 1,000, and these rarefied sample sums were used to quantify Shannon and Simpson α-diversity indices. Eight additional beetles (#SampleID 259, 280, 271, 187, 239, 175, 247, 151; Supplemental Table 4) had fewer than 1000 reads and were removed from only the α-diversity quantification. These eight individuals were retained in all other analyses. For analyses other than α-diversity, read counts were normalized to the total number of reads obtained for each individual sample (McMurdie and Holmes 2014). To calculate ß-diversity, only the top 30 most abundant OTUs for the three sub-groups of samples were used (*Acalymma* collected from different states; *Acalymma vittatum* collected within a season; and *Eucera (P*.*) pruinosa*). Distance matrices were calculated using unweighted Unifrac, and a PcoA of these distances were constructed using Phyloseq and custom scripts (McMurdie and Holmes 2013). A PERMANOVA adonis test on the unweighted Unifrac distance matrix was implemented in vegan (Dixon 2003). All plots were created with phyloseq (McMurdie and Holmes 2013) and ggplot2 (Wickham 2016).

## Results

### Sequencing Summary

In the United States, *Eucera (P*.*) pruinosa* samples were collected from cultivated *Cucurbita* plants in Pennsylvania and California. *Acalymma vittatum* and *A. trivittatum* were collected from cultivated plants in New Mexico, Arizona, Alabama, Iowa, Kentucky, Massachusetts and Vermont in the United States. *Acalymma trivittatum* were also collected from wild *Cucurbita* plants in Queretero, Mexico. At each location, all insects were sampled from the same agricultural field, or same wild plant population. In total, bacterial communities of 59 *Acalymma* beetles and 11 *Eucera pruinosa* bee individuals were sequenced (Figure 1a and 1b, Table 1). Sequencing the V3-V4 region of the 16S rRNA gene resulted in an average of 13,696 reads per sample (when grouped at 97% similarity, and after filtering out reads belonging to OTUs assigned to chloroplast, mitochondria and *Wolbachia*). There were 1,326 unique *de novo* OTUs overall, which were classified into 17 unique phyla and 149 unique families.

**Figure 1:**
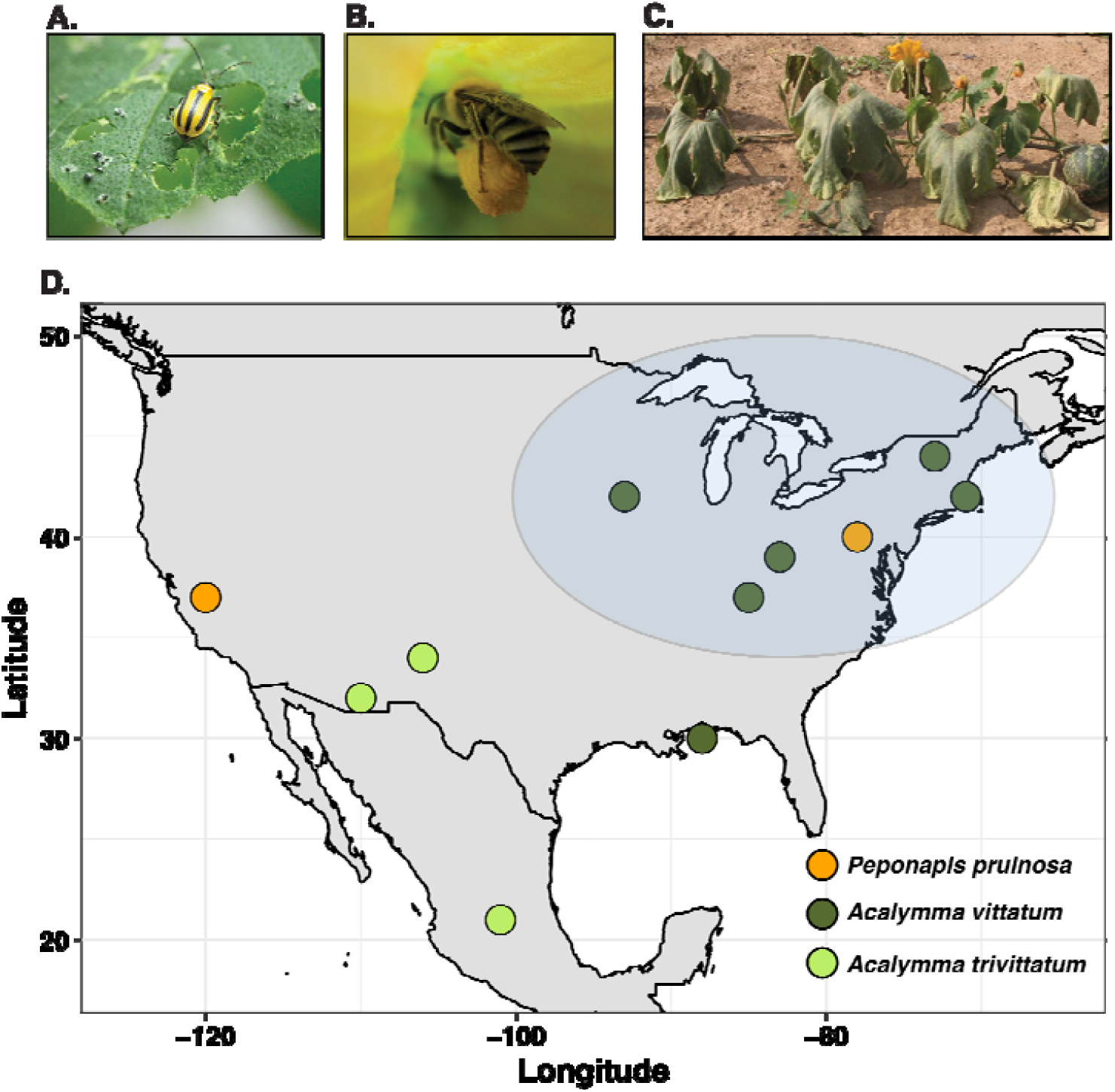
The study system. **A)** The Eastern striped cucumber beetle, *Acalymma vittatum* resting on a *Cucurbita pepo* leaf with heavy herbivory damage. This species is the most important herbivore of cucurbits in temperate Eastern North America, and is the predominant vector driving *Erwinia tracheiphila* transmission dynamics **B)** The Eastern squash bee, *Eucera (Peponapis) pruinosa*, gathering pollen and nectar from a male *Cucurbita pepo* flower **C)** A vine of a wilting, *Erwinia tracheiphila* infected wild gourd, *Cucurbita pepo* ssp. *texana*. **D)** The locations where bees and beetles sequenced in this study were collected. The blue oval approximates the region where *Erwinia tracheiphila* is an annual epidemic

### *Acalymma vittatum and* A. trivittatum *microbiome composition and geographic variation*

There were a total of 1,179 unique *de novo* OTUs present in the 59 sequenced *Acalymma* leaf beetle samples, and an average of 148 OTUs in each individual beetle. There are 16 OTUs that each comprise at least 1% of the total bacterial abundance, and collectively these 16 OTUs account for 83% of the total bacterial abundance across all the *Acalymma* individuals analyzed (Supplemental Table 1). These 16 OTUs belong to Gammaproteobacteria (50%; 8 of 16), Betaproteobacteria (12.5%; 2 of 16), Lactobacillales (12.5%; 2 of 16), Bacteroidetes (12.5%; 2 of 16) and Actinobacteria (6.25%; 1 of 16).

The taxonomic distribution of the 1,163 remaining rare OTUs (that are each present as less than 1% of the overall bacterial abundance) is similar to the taxonomic distribution of the 16 most common OTUs. Most of the rare OTUs are from Proteobacteria (44%; 514 out of 1163), followed by OTUs from Firmicutes (26%; 304 out of 1163), Actinobacteria (18.5%; 215 out of 1163), Chloroflexi (3.4%; 39 out of 1163) and Bacteroidetes (2.9%; 34 out of 1163). Other rare, low abundance OTUs are assigned to Acidobacteria, Verrucomicrobia, Plantomycetes, Mollicutes and several Archeal phyla (Funaro et al. 2011, Abdul Rahman et al. 2015, Alves et al. 2016). There was moderate statistical support for differences in the α-diversity (number and evenness) of OTUs in beetles collected in the different geographic locations (Kruskal-Wallis χ^2^=11.9, df = 6, *P =* 0.06), but there is no clear geographic factor underlying these differences (Figure 2). The bacterial community composition of beetles collected in the different states is qualitatively different overall (adonis PERMANOVA on unweighted UniFrac distances of the 30 most abundant OTUs; df = 6, *P* ≤ 0.01). Beetles collected from Iowa (USA) have the most homogeneous communities, while beetles collected from Arizona (USA) and Querétaro (MX) – both locations in the ancestral native range of *A. vittatum* – have the most variable composition.

**Figure 2:**
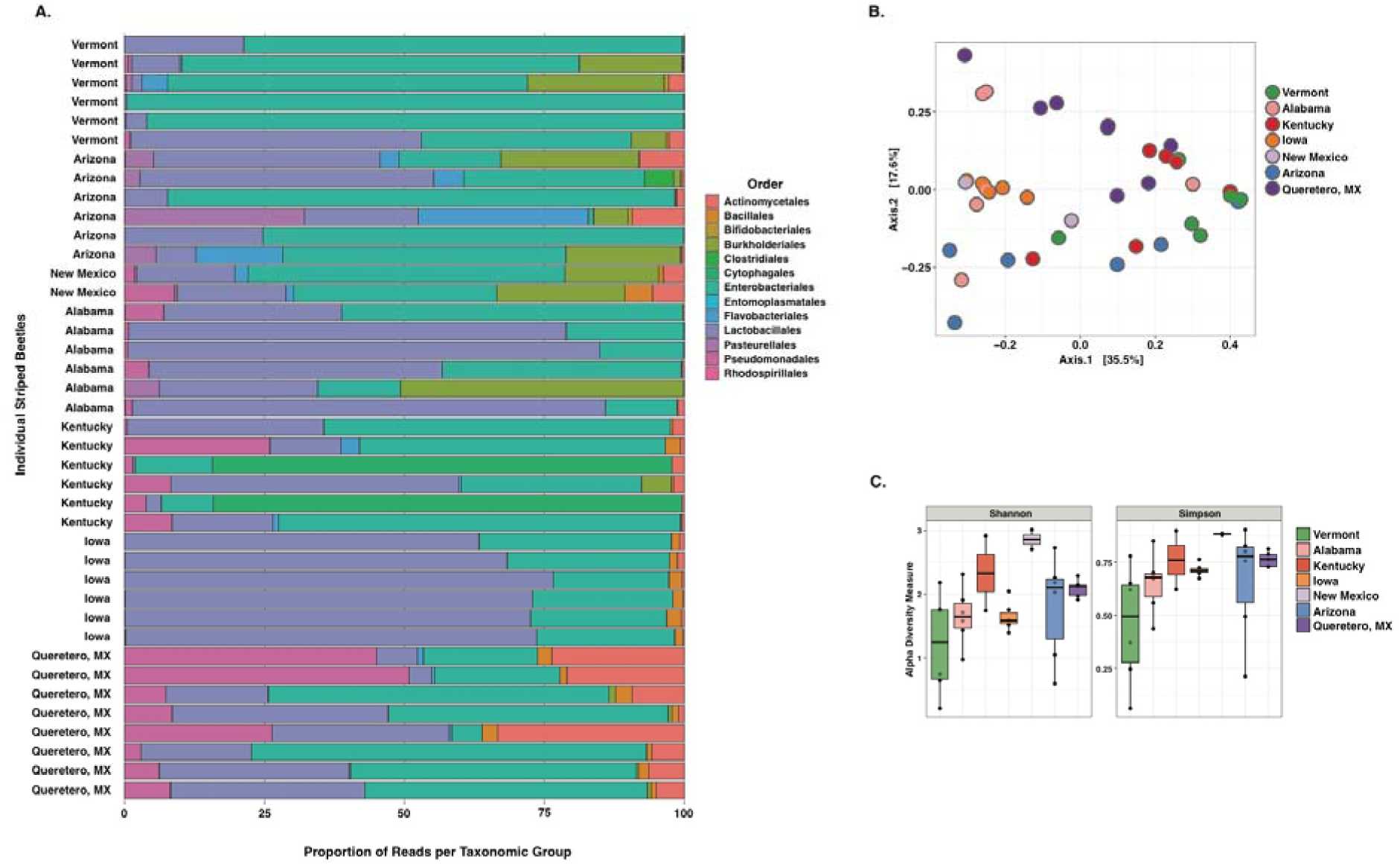
Summary of bacterial community composition of *Acalymma vittatum* and *trivittatum*. **A)** Bacterial community composition of individual *Acalymma vittatum* and *trivittatum* beetles. Each horizontal bar is the bacterial community composition in an individual beetle. The collection location of each sample is labeled on the Y-axis. Community composition is displayed by combining OTUs within each order, and the counts in each insect were normalized to the total number of reads in that individual and displayed as a percentage of the total community. **B)** Principle coordinates analysis (PCoA) of unweighted Unifrac distances, with each dot representing an individual insect. Insects are colored according to collection location. **C)** Shannon and Simpson α-diversity indices for the bacterial communities of *Acalymma vittatum* and *trivittatum* collected from different geographic locations (corresponding to Table 1 and Figure 1D). The number of reads per insect was rarefied to an even depth of 1000. Black dots are individual insects, and boxplots show the median and the quartile variation.

### *Within-season variation of* Acalymma vittatum *bacterial communities*

To quantify how seasonality affects the composition of beetle gut bacterial communities, adult *Acalymma vittatum* were collected from cultivated *Cucurbita* plants in Cambridge, Massachusetts (USA) soon after emerging from diapuase in the spring (June 7, 2016), from the second generation in the middle of the growing season (July 12, 2016), and from the diapausing generation towards the end of the growing season (August 14, 2016). There is high variability in α-diversity in individual *A. vittatum* collected throughout a single season, and no time point had significantly higher or lower α-diversity than the others (Kruskal-Wallis χ^2^= 0.61, df = 2, *P =* 0.74) (Figure 3). However, the composition of beetle bacterial communities does significantly change over the course of the growing season (adonis PERMANOVA on unweighted UniFrac distances of the 30 most abundant OTUs df = 2, *P* ≤ 0.01). Beetles collected soon after emerging from underground winter diapause (June 7, 2016) have higher abundances of Actinomycetales and Cytophagales – which are thought to predominantly be soil-dwelling species – than the other two time points. Beetles collected in the middle of the season (July 12, 2016), when many *Cucurbita* cultivars are flowering (and beetles are likely consuming protein-rich pollen and sugar-rich nectar (Samuelson 1994, Sasu et al. 2010a, Shapiro et al. 2012)) have the highest abundance of Lactobacillales. Bacterial communities of beetles collected on August 14 are dominated by OTUs assigned to Enterobacteriales and Pseudomonadales, which are likely derived from foliar tissue.

**Figure 3:**
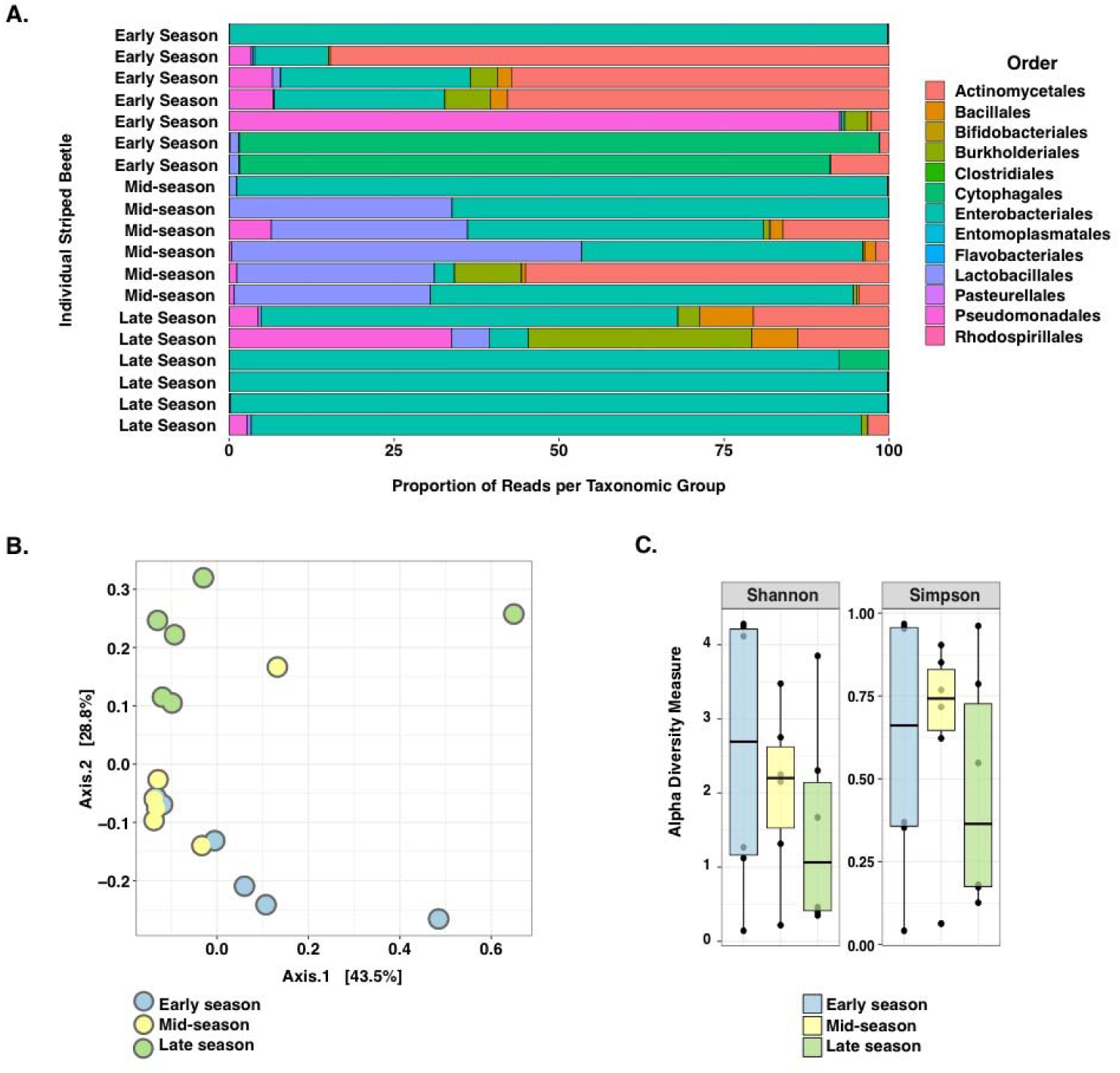
Seasonal variation of microbial communities from the same population of *Acalymma vittatum* in a Massachusetts. **A)** Bacterial community composition of individual *Acalymma vittatum* beetles collected in Cambridge, Massachusetts. Beetles were collected early in the season as adults emerged from winter diapause and cucurbits had just being planted (June 7 2016); in the middle of the season when commercial summer squash are blooming (July 12 2016); and at the end of the season as the second generation was nearing diapause (August 2016). The collection group each insect belongs to is labeled on the Y-axis. Community composition is displayed by combining OTUs within each order, and the counts in each insect were normalized to the total number of reads in that individual and displayed as a percentage of the total community. **B)** Principle coordinates analysis (PCoA) of unweighted Unifrac distances, with each dot representing an individual insect. Insects are colored according to date of collection. **C)** Shannon and Simpson α-diversity indices for the gut bacterial communities of *Acalymma vittatum* collected at three time points in a single growing season. The number of reads per insect was rarefied to an even depth of 1000. Black dots are individual insects, and boxplots show the median and the quartile variation.

### *Prevalence of* Erwinia tracheiphila *in* Acalymma vittatum *and* A. trivittatum *beetles*

*Erwinia tracheiphila* cannot be differentiated from closely related *Erwinia* and *Pantoea* strains with certainty from the sequences of V3-V4 rRNA gene fragments. To be conservative about estimating *E. tracheiphila* prevalence and abundance, all OTUs that were assigned to Enterobacteriaceae, and their taxonomic assignments were manually curated. Eight OTUs had at least a 98% sequence similarity to *E. tracheiphila* as the first or second BLASTN (Altschul et al. 1990) hit (using the NCBI rRNA reference database), and the prevalence and abundance of these OTUs were combined and all considered as putative *E. tracheiphila* (Supplemental File 3). Even using this permissive criteria that almost certainly over-estimates *E. tracheiphila* presence, only 33 out of the 59 *Acalymma* beetles harbor any of these putative *E. tracheiphila* OTUs (Table 2). When *E. tracheiphila* OTUs are detected in an individual beetle, they never sum to more than 1% of the total bacterial abundance within that insect (Table 2). Surprisingly, *E. tracheiphila* OTUs are detected not only in *Acalymma* beetles collected in temperate Easterns regions where this pathogen causes annual epidemics, but was also detected in a number of beetles collected from Tucson, AZ, USA and Querétero, MX – areas far from where *E. tracheiphila* is known to occur. This suggests there may be *Erwinia* strains or species closely related to *Erwinia tracheiphila* that may be common, non-pathogenic commensals of wild *Cucurbita* plants.

**Table 2:** Percentage of the total bacterial community in each individual insect that is *Erwinia tracheiphila*

| Individual Sample ID | Location | *Erwinia tracheiphila* geographic zone | Total Number of Bacterial Reads (top 20 OTUs) | *Total Number of Reads in OTUs near *Erwinia tracheiphila* | Percentage of total reads putatively *Erwinia tracheiphila* |
| --- | --- | --- | --- | --- | --- |
| 6 | Massachusetts (early) | Yes | 1544 | 0 | 0 |
| 203 | Massachusetts (early) | Yes | 387 | 0 | 0 |
| 215 | Massachusetts (early) | Yes | 13763 | 1 | 0.0073 |
| 239 | Massachusetts (early) | Yes | 623 | 0 | 0 |
| 251 | Massachusetts (early) | Yes | 7488 | 0 | 0 |
| 263 | Massachusetts (early) | Yes | 43316 | 18 | 0.0416 |
| 275 | Massachusetts (early) | Yes | 247 | 0 | 0 |
| 106 | Massachusetts (mid-season) | Yes | 10217 | 0 | 0 |
| 118 | Massachusetts (mid-season) | Yes | 2972 | 4 | 0.1346 |
| 155 | Massachusetts (mid-season) | Yes | 2990 | 2 | 0.0669 |
| 167 | Massachusetts (mid-season) | Yes | 1448 | 1 | 0.0691 |
| 179 | Massachusetts (mid-season) | Yes | 6280 | 1 | 0.0159 |
| 191 | Massachusetts (mid-season) | Yes | 28553 | 4 | 0.0140 |
| 90 | Massachusetts (late-season) | Yes | 732 | 0 | 0 |
| 105 | Massachusetts (late-season) | Yes | 1160 | 5 | 0.4310 |
| 117 | Massachusetts (late-season) | Yes | 19665 | 59 | 0.3000 |
| 129 | Massachusetts (late-season) | Yes | 14516 | 59 | 0.4064 |
| 141 | Massachusetts (late-season) | Yes | 1734 | 0 | 0 |
| 165 | Massachusetts (late-season) | Yes | 2765 | 13 | 0.4702 |
| 101 | Vermont | Yes | 5760 | 2 | 0.0347 |
| 138 | Vermont | Yes | 22983 | 14 | 0.0609 |
| 150 | Vermont | Yes | 7344 | 58 | 0.7898 |
| 162 | Vermont | Yes | 6830 | 0 | 0 |
| 174 | Vermont | Yes | 12054 | 27 | 0.2240 |
| 186 | Vermont | Yes | 17541 | 6 | 0.0342 |
| 260 | Alabama | No | 1041 | 0 | 0 |
| 248 | Alabama | No | 2638 | 0 | 0 |
| 236 | Alabama | No | 2518 | 0 | 0 |
| 224 | Alabama | No | 4531 | 0 | 0 |
| 212 | Alabama | No | 13223 | 2 | 0.0151 |
| 200 | Alabama | No | 4651 | 7 | 0.1505 |
| 268 | Kentucky | Yes | 1080 | 1 | 0.0926 |
| 280 | Kentucky | Yes | 258 | 0 | 0 |
| 283 | Kentucky | Yes | 1630 | 0 | 0 |
| 271 | Kentucky | Yes | 455 | 0 | 0 |
| 259 | Kentucky | Yes | 138 | 0 | 0 |
| 247 | Kentucky | Yes | 641 | 0 | 0 |
| 190 | Iowa | Yes | 9058 | 2 | 0.0221 |
| 272 | Iowa | Yes | 10720 | 0 | 0 |
| 166 | Iowa | Yes | 3849 | 0 | 0 |
| 154 | Iowa | Yes | 2544 | 0 | 0 |
| 142 | Iowa | Yes | 6035 | 0 | 0 |
| 130 | Iowa | Yes | 4014 | 1 | 0.0249 |
| 257 | New_Mexico | No | 2525 | 1 | 0.0396 |
| 269 | New_Mexico | No | 1856 | 0 | 0 |
| 216 | Arizona | No | 27221 | 1 | 0.0037 |
| 204 | Arizona | No | 36866 | 19 | 0.0515 |
| 40 | Arizona | No | 56257 | 0 | 0 |
| 28 | Arizona | No | 48542 | 5 | 0.0103 |
| 16 | Arizona | No | 51246 | 67 | 0.1307 |
| 4 | Arizona | No | 57625 | 10 | 0.0174 |
| 187 | Querétaro MX | No | 557 | 0 | 0 |
| 78 | Querétaro MX | No | 14615 | 16 | 0.1095 |
| 175 | Querétaro MX | No | 644 | 2 | 0.3106 |
| 163 | Querétaro MX | No | 14801 | 15 | 0.1013 |
| 151 | Querétaro MX | No | 719 | 0 | 0 |
| 139 | Querétaro MX | No | 1469 | 1 | 0.0681 |
| 127 | Querétaro MX | No | 8398 | 58 | 0.6906 |
| 66 | Querétaro MX | No | 52178 | 483 | 0.9257 |

\*The number of reads in all individual putative *Erwinia tracheiphila* OTUs were summed

### Eucera (P.) pruinosa *microbiome composition*

Overall, there is a total of 312 distinct *de novo* OTUs present in at least one of the 11 sequenced *Eucera (P*.*) pruinosa* samples (Table 1, Supplemental Table 2). There is an average of 93 OTUs in each individual bee collected in California, while there is an average of only 30 OTUs in each bee collected in Pennsylvania. There are 14 OTUs that comprise at least 1% of the total bacterial abundance, and these 14 OTUs collectively sum to 89.6% of the total bacterial abundance in the *Eucera* bee samples. Of the 14 common OTUs, 6 are assigned to Lactobacillales (Bacilli), 5 are assigned to Enterobacteriales and Pseudomonadales (Gammaproteobacteria), 1 is assigned to Actinobacteria and 1 to Entoplasmatales (Mollicutes). The most abundant OTU in *E. (P*.*) pruinosa* (*de novo* 415107) is assigned to *Acinetobacter* (Gammaproteobacteriaceae), a genus with many species commonly associated with floral nectar (Álvarez-Pérez et al. 2013, McFrederick and Rehan 2016). The Lactobacillales OTUs are likely involved in breakdown of pollen, and/or fermentation of nectar carbohydrates (McFrederick et al. 2012, McFrederick et al. 2018, Vuong and McFrederick 2019). Pollen is very high in fructose, and one of the 14 common OTUs is assigned to *Fructobacillus* (*de novo* 382787), which may also function to breakdown sugar in pollen and nectar (Endo and Salminen 2013). The same *Fructobacillus* OTU is occasionally present in *Acalymma*, but at extremely low abundance Supplemental Table 1). One of the 14 common *E. (P*.*) pruinosa* OTUs is assigned to *Spiroplasma* sp. nr. *apis* (Entoplasmatales) (*de novo* 410890), and is common in bees collected in both CA and PA. Some Entoplasmatales species are ancient commensals of insects (Funaro et al. 2011), while other species are emerging as pathogens of wild and managed bees (Ravoet et al. 2014). Of the 299 rare OTUs that comprise the remaining 11% of bacterial abundance, most OTUs are assigned to Gammaproteobacteria (43%; 129 out of 299), followed by Lactobacillales (37%; 111 out of 299), Actinobacteria (6%; 19 out of 299) and Entomoplasmatales (3.7%; 11 out of 299) (Supplemental Table 2). *E. (P*.*) pruinosa* collected in California have marginally higher average α-diversity (Kruskal-Walls χ^2^= 1.75, df = 1, *P =* 0.18) (Figure 4), and the bacterial communities from bees sampled in CA vs. PA are marginally qualitatively different (adonis PERMANOVA on unweighted UniFrac distances of the 30 most abundant OTUs df = 2, *P =* 0.09).

**Figure 4:**
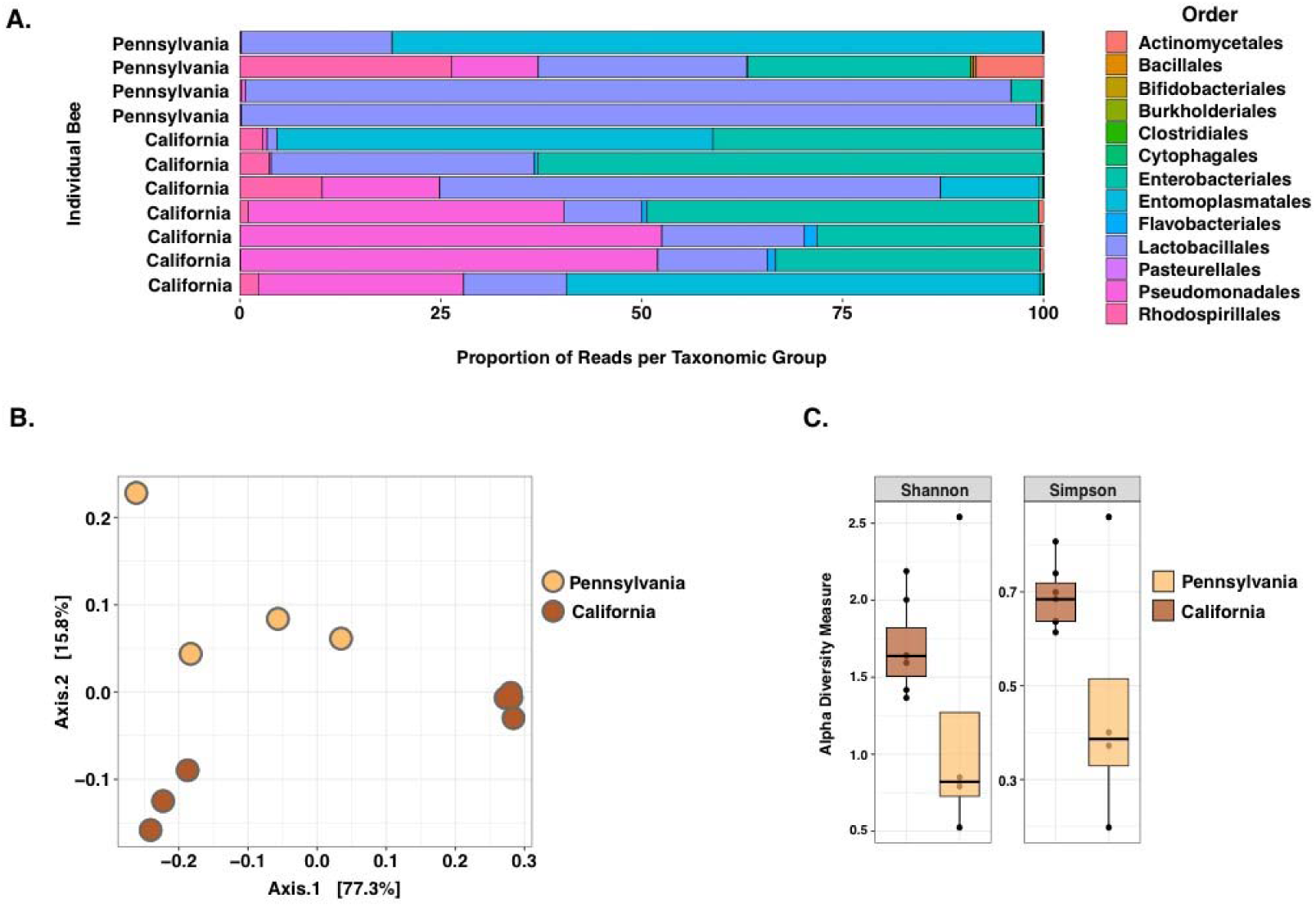
Geographic variation in *Eucera (Peponapis) pruinosa* bacterial communities. **A)** Bacterial community composition of individual *Eucera (P*.*) pruinosa* bees collected in California or Pennsylvania. The collection location of each sample is labeled on the Y-axis. Community composition is displayed by combining OTUs within each order, and the counts in each insect were normalized to the total number of reads in that individual and displayed as a percentage of the total community. **B)** Principle coordinates analysis (PCoA) of unweighted Unifrac distances, with each dot representing an individual insect. Insects are colored according to collection location. **C)** Shannon and Simpson α-diversity indices for the gut bacterial communities of *Eucera (P*.*) pruinosa* collected in California vs. Pennsylvania. The number of reads per insect was rarefied to an even depth of 1000. Black dots are individual insects, and boxplots show the median and the quartile variation.

### *Comparison of microbial communities in* Acalymma *and* Eucera (P.) pruinosa

Despite the high total number of *de novo* bacterial OTUs, six of the top 7 most abundant OTUs in *Acalymma* beetles are also among the 14 most abundant OTUs in *Eucera* bees (Supplemental Table 1, Supplemental Table 2, (Figure 5) (Herrera et al. 2009, Fridman et al. 2012). As expected, the taxonomic assignments of the OTUs found in *E. pruinosa* reflect microbes that are likely well-adapted to aid in digestion of the floral food sources. Surprisingly, many of these same common OTUs are found in *Acalymma*, suggesting that floral pollen and nectar may be an unappreciated nutritional source for these beetles, which are generally thought of as leaf herbivores due to the extensive foliar economic damage they inflict (Baker and Hurd Jr 1968, Samuelson 1994, Shapiro et al. 2012). A *Klebsiella* spp. OTU (*de novo* 423191) is the most abundant overall in *Acalymma* beetles, and the second most abundant in bees. Two Lactobacillales OTUs (*de novo* 432933 and *de novo* 222965), an Enterobacteriaeae (*de novo* 264045), an *Acinetobacter* (*de novo* 213309), and an Actinomycetales (*de novo* 211172) are also among the most common OTUs in both bees and beetles (Supplemental Table 4). This suggests these insects are exposed to these same OTUs through their shared floral food sources, and that these common OTUs may perform similar digestive functions related to breakdown of sugars (and perhaps proteins and fats) present in *Cucurbita* pollen and nectar.

**Figure 5:**
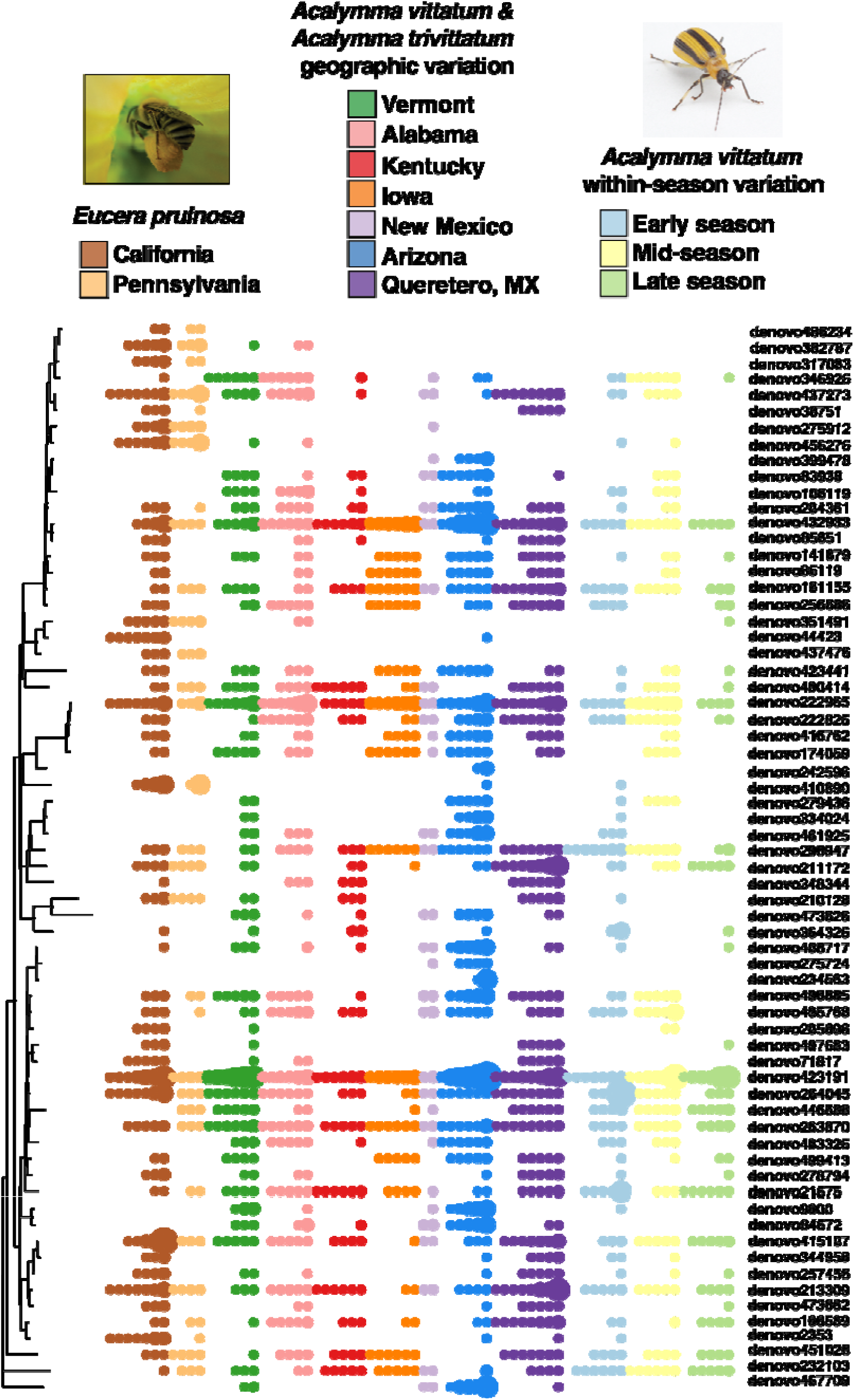
Prevalence and abundance of the most common bacterial OTUs in *Acalymma* beetles and *Eucera* (*Peponapis*) bees. The tree shows the phylogenetic distance relationships of the 50 most abundant OTUs in all *Acalymma* beetles and the 40 most abundant OTUs in the *Eucera (P*.*) pruinosa* bees. Each tip of the tree is an individual OTU, and corresponds to taxonomic assignments provided in Supplemental Table 4 Each colored dot represents the occurrence of that OTU in an individual insect, and the size of the dot corresponds to the relative abundance of that OTU in that individual.

### *Detection of* Wolbachia *in* Acalymma

Ten percent of *Acalymma vittatum* (6 out of 59 individuals) were infected with *Wolbachia*. This is consistent with a previous study that found a 5% *Wolbachia* infection rate of field-collected *A. vittatum* (Clark et al. 2001). All *Wolbachia-*positive beetles were collected from temperate Eastern North America (Massachusetts, Kentucky and Vermont) (Supplemental Figure 1). No *Acalymma* collected outside of the *E. tracheiphila* endemic area carried *Wolbachia*, and none of the *E. (P*.*) pruinosa* were positive for *Wolbachia*. The presence of *Wolbachia* in few beetles collected from within their newly expanded geographic range in temperate Eastern North America may be due to a recent shift of this reproductive parasite into *A. vittatum* as a host species. A better understanding of the effects of harboring *Wolbachia* on *A. vittatum* reproduction and population dynamics may be useful for developing sustainable vector control strategies (Clark et al. 2001, Kajtoch and Kotásková 2018).

## Discussion

Here, we find that field-collected *Acalymma vittatum* and *trivittatum* beetles, and to a lesser extent *Eucera (P*.*) pruinosa* bees are exposed to diverse bacteria during normal foraging. Squash bees and beetles have co-evolved to specialize on *Cucurbita*, and are both attracted over long distances to the large, yellow, highly scented flowers that secrete substantial quantities of sugary nectar, and produce large amounts of protein rich pollen (Metcalf and Lampman 1991, Mena Granero et al. 2005, Shapiro et al. 2012). *Cucurbita* flowers, when present, are a foraging site that likely also serves as a shared source of microbial exposure for both *Acalymma* and *Eucera* (*Peponapis*) (Baker and Hurd Jr 1968, Samuelson 1994, Salzman et al. 2018). Consumption of floral resources with the same nutrient and carbohydrate composition by both beetles and bees also likely drives the need for microbial associates that have the same metabolic functions for plant matter digestion.

The trend towards higher α-diversity in bees from California may be an artifact from the low sample sizes of bees sequenced in this study. Alternatively, this trend may reflect that in their native range, pollinators are exposed to a higher diversity of microbes while foraging on multiple co-occuring species of *Cucurbita*, and on both wild and cultivated genotypes. Populations of cultivated crops that are grown in homogeneous, human-managed landscapes have reduced genetic diversity compared to populations of wild relatives, and may also host microbial communities with lower diversity – measured as either 16S OTU richness, or total physiological potential of the microbial community. The susceptibility of crop plants grown in homogeneous, human managed agro-ecosystems to invasion by novel microbial pathogens is a well-documented and continuing risk (McDonald and Stukenbrock 2016, Shapiro et al. 2016, Shapiro et al. 2018b). How these homogeneous landscapes potentially also affect pollinator health through alterations in microbial ecology, perhaps by creating landscapes with lower microbial functional diversity, or increasing opportunities where wild and managed pollinators can exchange pathogens, remains unknown (Colla et al. 2006, Burke et al. 2011, Fürst et al. 2014, López-Uribe et al. 2016, McDonald and Stukenbrock 2016, Shapiro et al. 2016, Pérez-Jaramillo et al. 2017, Shapiro et al. 2018b, SA et al. 2019, Yu et al. 2019).

The high number of rare, low abundance OTUs in *Acalymma* herbivores is consistent with accumulating evidence that the local environment – *ie*, the variable and diverse microbial communities that colonize plants or the surrounding soil – are the most significant source of microbial exposure for most non-social and non-sap feeding insects (Moran 2003, Kwong and Moran 2016, Hammer et al. 2017, McFrederick et al. 2017, Hammer et al. 2019, Hannula et al. 2019). The lower average number of OTUs per insect in bees compared to beetles may reflect that bees exclusively feed on *Cucurbita* floral resources, while *Acalymma* beetles are often found feeding on other introduced cucurbit crop plants (most notably muskmelon and cucumbers, both *Cucumis* spp.), consume both foliage and floral structures, and can occasionally be found feeding on pollen from distantly related plant species (Gould 1944). *Acalymma* are highly mobile, and move frequently between foliage and flowers of different host plants and the surrounding soil during normal foraging, feeding and oviposition. The large geographic area from which beetles were sampled may also contribute to the high number of rare OTUs. Overall, the composition and diversity patterns reflect that *Acalymma* beetles are transiently exposed to diverse bacteria that do not persistently colonize them as hosts (Hammer et al. 2017, Hammer et al. 2019).

A diminishingly small proportion of the gut bacterial community in field-collected *A. vittatum* are OTUs that could be *Erwinia tracheiphila*. The low abundance of *E. tracheiphila* OTUs in beetle vectors is consistent with laboratory experiments to quantify *E. tracheiphila* in beeltes, and field studies to quantify transmission rates. In controlled laboratory experiments, populations of *E. tracheiphila* in infective vectors or frass were too low to be detected by standard PCR (Mitchell and Hanks 2009) but can be reliably detected by more sensitive fluorescent probe-based qPCR assays (Shapiro et al. 2014). In field trials conducted at Rock Springs Experimental Station in Central Pennsylvania (USA), almost all *Cucurbita* plants were exposed to *E. tracheiphila* every morning by beetles that gather to feed and mate in flowers (Sasu et al. 2010a). Despite this daily exposure to *E. tracheiphila* deposited by vectors, less than half of the experimental plants developed an active *E. tracheiphila* infection (Sasu et al. 2009, Sasu et al. 2010b). Genotypes of *Cucurbita* that produce more flowers, and more volatiles per flower – and therefore attract more foraging beetle vectors – are also associated with a significantly higher rate of *E. tracheiphila* infection in *Cucurbita pepo* (Sasu et al. 2009, Shapiro et al. 2012). This suggests that individual vectors deposit few *E. tracheiphila* cells during each feeding exposure, and that cumulative exposure to many infective vectors is more important than exposure to a single vector (Yao 1996).

The composition, function and variation of microbial communities that associate with most wild, unmanaged insect species is almost completely unknown. This study is the first step towards understanding factors underlying variation in the composition of bacterial communities that associate with the ecologically and economically important insect species that have co-evolved with *Cucurbita*. These results provide an important baseline for guiding future experimental studies to understand factors driving variation in bacterial community composition and functions. A more comprehensive understanding of which microbial strains and communities colonize *Eucera* and *Acalymma* across the diverse geographic they inhabit, and empirically testing the functions of individual isolates and more complex communities for the host insects will be important for developing effective vector controls and protecting pollinator health.

## Acknowledgements

LRS was supported by NSF postdoctoral fellowship DBI-1202736 and a grant from the North Carolina State Plant Soil Microbe Collaborative Consortium; JR was supported by Fundación Mexico en Harvard, and CONACYT grant 237414. We thank Roberto Kolter for experimental support and advice, Nick Sloff for Figure 1A beetle image, and Clubes de Ciencias Mexico, Ricardo Bessin, Salvador Montes, the University of Vermont Horticultural Farm and the gardeners at the Christian Herter Community Garden for assistance collecting beetles; the Harvard Odyssey high performance computational cluster for computing resources and Harvard Odyssey staff for compuational support. Mention of trade names or commercial products in this publication is solely for the purpose of providing specific information and does not imply recommendation or endorsement by the U.S. Department of Agriculture. Any opinions, findings, conclusion, or recommendations expressed in this publication are those of the author(s) and do not necessarily reflect the views of the U.S. Department of Agriculture.

---

**Supplemental Figure 1:** *Wolbachia* occurrence in *Acalymma* beetles.

**Supplemental Table 1:** Taxonomic assignments of OTUs, the number of individuals in which that OTU occurs, and the percentage of each OTU in the cumulative community in *Acalymma vittatum* and *trivittatum* beetles.

**Supplemental Table 2:** Taxonomic assignments of OTUs, the number of individuals in which that OTU occurs, and the percentage of each OTU in the cumulative community in *Eucera (P*.*) pruinosa* squash bees.

**Supplemental Table 3:** Manual curation of OTUs assigned to Enterobacteriaceae to assess which are most likely to be *Erwinia tracheiphila*.

**Supplemental Table 4:** Manual curation of taxonomic assignments of the dominant OTUs in *Acalymma* and *Eucera (Peponapis)*.

**Supplemental Table 5:** Primers, barcodes, and individual #SampleIDs for the insects analyzed in this study.

